# IL-33 controls IL-22-dependent antibacterial defense by modulating the microbiota

**DOI:** 10.1101/2023.07.19.549679

**Authors:** Ivo Röwekamp, Laura Maschirow, Anne Rabes, Facundo Fiocca Vernengo, Gitta Anne Heinz, Mir-Farzin Mashreghi, Sandra Caesar, Miha Milek, Anna Carolina Fagundes Fonseca, Sandra-Maria Wienhold, Geraldine Nouailles, Ling Yao, Dunja Bruder, Julia D. Boehme, Monika Puzianowska-Kuznicka, Dieter Beule, Martin Witzenrath, CAPNETZ Study Group, Max Löhning, Markus M. Heimesaat, Christoph S.N. Klose, Andreas Diefenbach, Bastian Opitz

## Abstract

IL-22 plays a critical role in defending against mucosal infections, but how IL-22 production is regulated is incompletely understood. Here, we show that mice lacking IL-33 or its receptor ST2 (IL-1RL1) were more resistant to *Streptococcus pneumoniae* lung infection than wild-type animals, and that single nucleotide polymorphisms in *IL33* and *IL1RL1* were associated with pneumococcal pneumonia in humans. The effect of IL-33 on *S. pneumoniae* infection was mediated by negative regulation of IL-22 production in innate lymphoid cells (ILCs), but independent of ILC2s as well as IL-4 and IL-13 signaling. Moreover, IL-33’s influence on antibacterial defense was dependent on housing conditions of the mice, and mediated by the modulatory effect of IL-33 on the microbiota. Collectively, we provide insight into the bidirectional crosstalk between the innate immune system and the microbiota. We identify a mechanism, dependent on both genetic and environmental factors, that impacts the efficacy of antibacterial immune defense and thus susceptibility to pneumonia.

**SIGNIFICANCE STATEMENT:** Lower respiratory tract infections are the fifth leading cause of death. Here, we describe a mechanism influenced by genetic and environmental factors that affects the efficacy of pulmonary antibacterial immune responses. We show that IL-33 controls antibacterial defense by regulating the production of IL-22, a cytokine with known functions in antimicrobial immunity in lungs. The effect of IL-33 on IL-22-dependent defense was influenced by the hygienic status of the mice and mediated by IL-33’s modulatory effect on the animal microbiota. In addition, genetic variation in genes involved in IL-33 signaling was associated with bacterial pneumonia in humans. Our findings may be important for our understanding of the factors influencing predisposition to lower respiratory tract infections.

## INTRODUCTION

Lower respiratory tract infections are the most common infectious cause of death and the fifth leading cause of mortality overall (1). *Streptococcus pneumoniae* is the most frequently found pathogen in community-acquired pneumonia (CAP) (2, 3). The extracellular bacterium often asymptomaticly colonizes the mucosal surfaces of the upper respiratory tract. Pneumonia only develops if pneumococci get access to the distal airways by aspiration and if they resist immediate elimination by the immune system (4, 5). The factors influencing the efficacy of antibacterial immune defense pathways in the lungs and thus determining susceptibility to pneumonia are, however, incompletely understood.

Antimicrobial immune responses to infections are initiated after sensing of microbial molecules by pattern recognition receptors (6, 7). This initiates release of chemokines and cytokines such as TNF-α, IL-1β and IL-23, some of which induce subsequent production of additional cytokines including IFNγ and IL-22 by lymphoid cells (8). IL-22, which is primarily produced by type 3 innate lymphoid cells (ILCs), γδ T cells and Th17 cells, is a mediator of immune defense against extracellular pathogens at mucosal organs (9-13). During pulmonary infections, IL-22 induces expression of antimicrobial peptides, tight junction proteins and complement factors by lung epithelial cells and hepatocytes, respectively (14-17).

Alarmins such as HMGB1, IL-33, ATP and uric acid are best known for their role as mediators of inflammation in response to sterile tissue damage (18). They are generally defined as endogenous danger molecules, which are normally hidden inside of host cells and only released upon cellular damage or stress. IL-33, for example, is stored in the nucleus of various stroma cells, and is best known for its capacity to drive type 2 inflammation after release and binding to its transmembrane receptor ST2 (encoded by *IL1RL1*) (19). ST2 is expressed by e.g. Th2 cells, ILC2s, and regulatory T cells but also by activated Th1 and CD8^+^ cytotoxic T cells (20, 21). Moreover, a soluble form of ST2 (sST2) is generated by alternative polyadenylation, which lacks the transmembrane domain and binds IL-33 as a decoy receptor to prevent it from activating ST2 signaling (21, 22).

In our study, we initially aimed for a better understanding of the role of alarmins in regulating innate immune responses during bacterial infections. We demonstrate that IL-33 controls immune defense against *S. pneumoniae* in the lungs by influencing IL-22 production. Unexpectedly, however, we found that this mechanism was independent of ILC2s, IL-4 and IL-13. Instead, microbiota characterization by shotgun sequencing as well as functional cohousing and microbiota depletion experiments show that IL-33’s influence on *S. pneumoniae* infection was mediated by its modulatory effect on the microbiota.

## RESULTS

### IL-33 enhances susceptibilty to S. pneumoniae-induced pneumonia

To better understand how the immune response to *S. pneumoniae* is regulated, we first investigated if alarmins were released during infection. We detected increased levels of uric acid, ATP, and IL-33 in the supernatants of macrophages, alveolar epithelial cells and/or lung tissue explants upon infection with *S. pneumoniae* in vitro (Fig. S1A-C). Next, we tested if these alarmins affect pneumococcal pneumonia in vivo. As assessed 48 h after intranasal infection, neither degradation of uric acid or ATP, nor inhibition of or deficiency in nucleotide receptors influenced bacterial loads (Fig. S1D-G). In contrast, lack of IL-33 or ST2 rendered mice more resistant to *S. pneumoniae*, as indicated by improved bacterial control (Fig. 1A, B, D, E) and reduced body temperature decrease (Fig. 1C, F). Bone marrow chimera experiments, using WT or *Il33*^-/-^ mice as either donors or recipients of bone marrow transplants, indicated that the detrimental effect of IL-33 during pneumococcal infection depended on its expression in recipient animals rather than the transplanted bone marrow (Fig. 1G, H). In addition, we analyzed single nucleotide polymorphisms (SNPs) previously associated with type 2 inflammation and altered sST2 or IL-33 levels (22-27) in 238 pneumococcal pneumonia patients and 238 age- and sex-matched controls. We found that SNP alleles previously linked to lower levels of the decoy receptor sST2 (22, 23), and by trend a SNP allele recently associated with higher IL-33 levels (24), were more frequent in the pneumonia patients as compared to the controls (Fig. S2A-D). Overall, our results demonstrate that IL-33 influences susceptibility to pneumococcal pneumonia in mice and possibly in humans.

**Figure 1.**
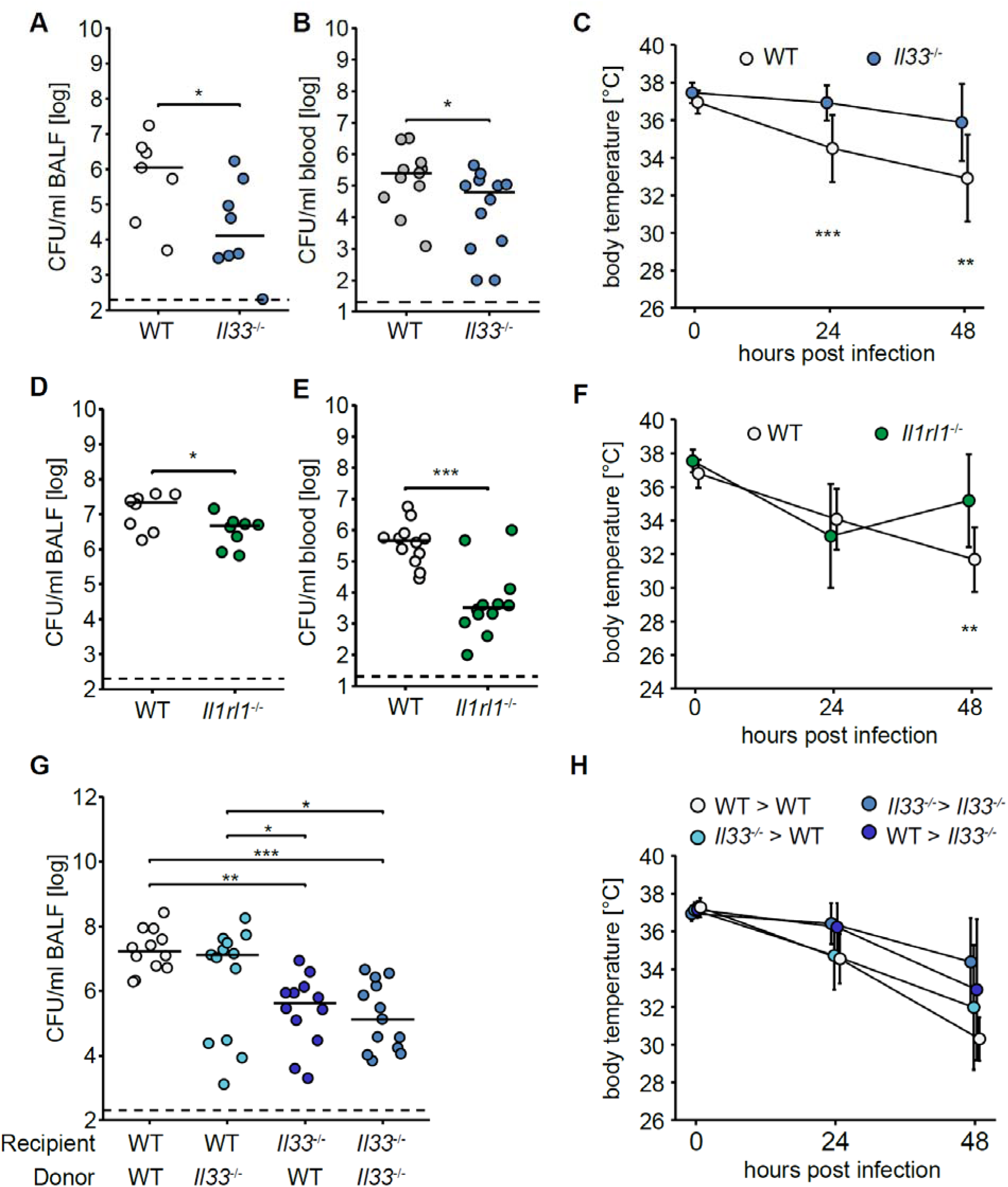
IL-33 deficiency enhances resistance of mice to pneumococcal pneumonia. WT and *Il33*^-/-^ were intranasally infected with 5×10^6^ CFU/mouse of *S. pneumoniae* and bacterial loads in BALF (**A**) and blood (**B**) were determined 48 h post infection, and temperature was measured at the indicated time points (**C**). (**D - F**) WT and *Il1rl1*^-/-^ were infected and bacterial loads in BALF (**D**) and blood (**E**) were determined 48 h post infection, and temperature was measured (**F**). (**G, H**) Bone marrow chimeras were infected and bacterial loads in BALF were determined 48 h post infection (**G**), and temperature was measured (**H**). (**A, B, D, E,G**) Data are shown as individual data points. Lines represent median and dashed line the lower detection limit. Wilcoxon rank sum test or Kruskal-Wallis followed by Dunn’s posthoc test were used if more than two groups were compared,. (**C, F, H**) Data are shown as mean ± SD, Kruskal-Wallis followed by Dunn’s posthoc test; * = p < 0.05, ** = p < 0.01, *** = p < 0.001.

### Lack of ILC2s does not enhance susceptibility to S. pneumoniae

To characterize the mechanism underlying the enhanced resistance of IL-33-deficient mice to *S. pneumoniae*, we first examined their inflammatory immune response. We did not observe any influence of IL-33 deficiency on levels of chemokines and various cytokines in bronchial alveolar lavage fluid (BALF) (Fig. S3A), nor did we observe an effect on numbers and proportions of alveolar macrophages (AMs), polymorphonuclear neutrophils (PMNs) and inflammatory monocytes (Fig. S3B-D). Next, we performed single cell RNA sequencing (scRNAseq) of pulmonary cells from *S. pneumoniae*-infected mice. We enriched the total lung cells (3/4) with FACS-sorted lineage^-^ CD90.2^+^ CD127^+^ cells (1/4) to also enable analysis of scarcer ILCs. Cellular populations were visualized by Uniform Manifold Approximation And Projection (UMAP) (Fig. S4A) and classified based on signature genes (Fig. S4B). We found *IL1RL1* to be primarily expressed by ILC2s (Fig. 2A). In addition, we defined clusters for AMs, PMNs and monocytes based on unbiased clustering, but did not observe differences in their relative abundance between WT and *Il33*^-/-^ mice (Fig. S4C-K).

**Figure 2.**
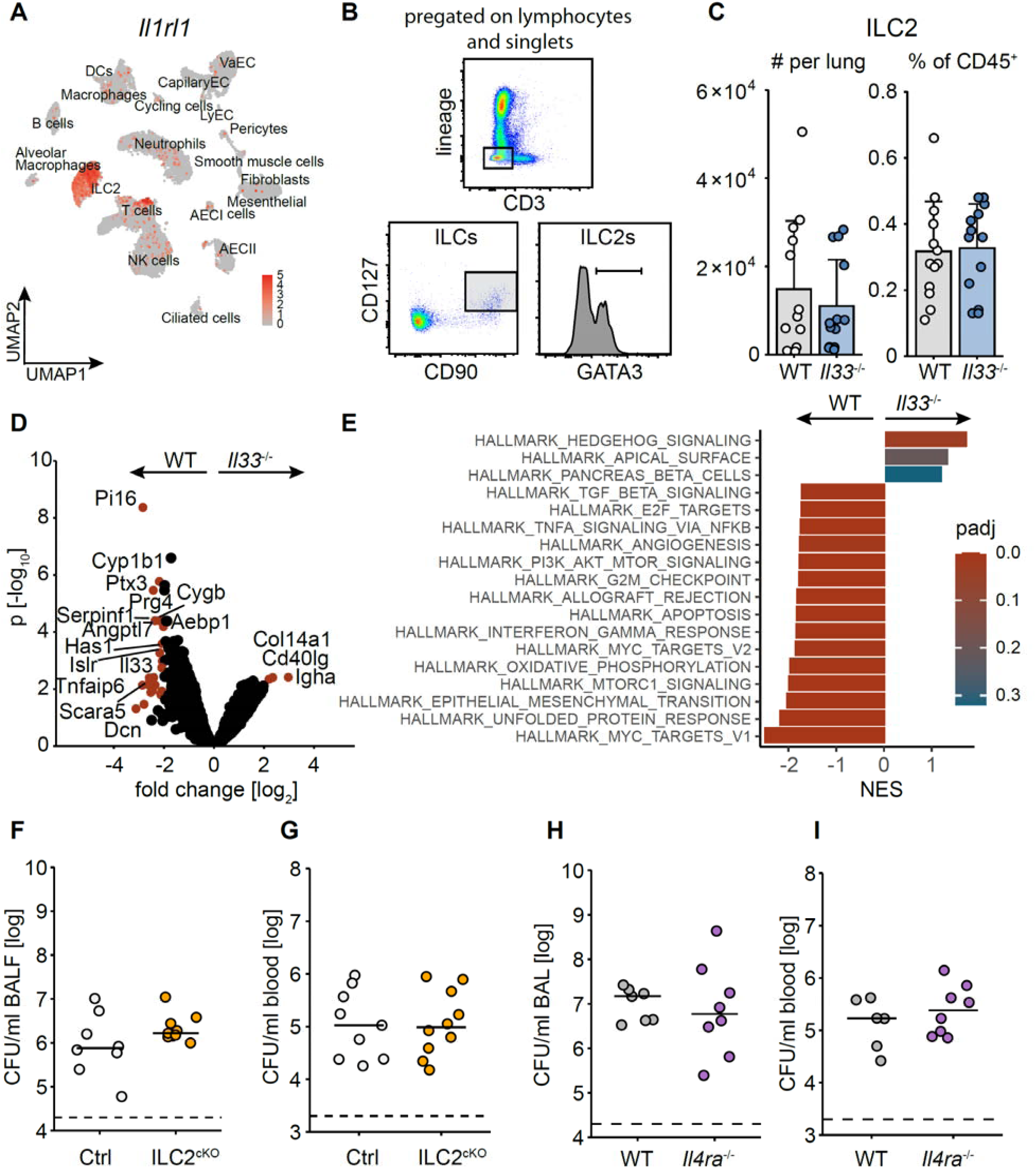
ILC2 as well as IL-4 and IL-13 signaling do not regulate early antibacterial defense during pneumococcal pneumonia. (**A**) WT and *Il33*^-/-^ mice were infected with *S. pneumoniae*, sacrificed after 18 h and pulmonary cells were analysed by scRNAseq (pulmonary cells were pooled from n = 4 per group). Normalized expression levels of *Il1rl1* were plotted in an UMAP embedding. (**B**) Gating strategy to quantify ILC2s by FACS. (**C**) Numbers and frequencies of pulmonary Lin^-^ CD90^-^ CD127^-^ GATA3^+^ ILC2s from lungs of mice 18 h after infection. Bars represent mean + SD (**D, E**) Lungs were digested at 12 h post infection and sorted for ILC2s, RNA was isolated and quantified using bulk RNA-sequencing. (**D**) Volcano plot displaying differentially expressed genes (DEGs) in ILC2s of WT and *Il33*^-/-^animals. Significant DEGs with a log2FC threshold of ± 2 and adjusted P value <0.1 are indicated in red. (**E**) GSEA analysis utilizing HALLMARK collection, normalized enrichment score (NES) and adjusted p-values (padj) are indicated. (**F, G**) *Nmur1*^iCre-eGFP^*Id2*^fl/fl^ mice (ILC2^cKO^) and controls were infected and bacterial loads in BALF and blood were assessed 48 hours post infection (**H, I**) *Il4ra*^-/-^ and control mice were infected and bacterial loads in BALF and blood were assessed at 48 hours post infection. Data are shown as individual data points. Lines represent median and dashed line the lower detection limit.

Since the IL-33 receptor was mainly expressed by ILC2s, we next compared their numbers and proportions in WT and *Il33*^-/-^ mice, without identifying differences (Fig. 2B, C). Next, we investigated their gene expression by bulkRNAseq. Several genes including, for example, *Pi16*, *Cyp1b1*, and *Tnfaip6* and gene sets such as “HALLMARK_MYC_TARGETS” showed decreased expression in *Il33*^-/-^ as compared to WT mice during *S. pneumoniae* infection (Fig. 2D, E). Early control of *S. pneumoniae* infection was, however, unchanged in mice specifically lacking ILC2s (*Nmur1^iCre-eGFP^Id2^fl/fl^*) (28, 29) or in animals deficient in IL-4 and IL-13 signaling (*Il4ra*^-/-^) (Fig. 2F-I). Thus, unlike IL-33 and ST2, ILC2s or the type 2 inflammation-associated cytokines IL-4 and IL-13 do not regulate early antibacterial defense against *S. pneumoniae* during lung infection.

### Protective effect of IL-33 deficiency in pneumococcal infection is mediated by enhanced IL-22 production

IL-22 is an important mediator of antibacterial immune defense at mucosal barriers such as the lungs (30, 31). We noticed enhanced IL-22 production by *Il33*^-/-^ animals as compared to WT controls upon infection with *S. pneumoniae* (Fig. 3A). In line with this observation, we found a significant increase in the proportions and numbers of IL-22-producing ILCs in the lungs of infected *Il33*^-/-^ mice, which were ST2-negative and most likely represented ILC3s, but we did not observe differences in IL-22 production by αβ and γδ T cells (Fig. 3B, C, D). In order to test if the enhanced production of IL-22 was responsible for the increased resistance of *Il33*^-/-^ mice to *S. pneumoniae* infection, we generated *Il33*^-/-^*Il22*^-/-^ double knockout mice and compared them with *IL33*^-/-^, *Il22*^-/-^ and WT animals. We observed that the bacterial loads in lungs and blood of *Il33*^-/-^ *Il22*^-/-^ mice were largely similar compared to the ones in WT mice and significantly higher than in *Il33*^-/-^ mice (Fig. 3E, F). These results indicate that increased production of IL-22 is responsible for the enhanced resistance of IL-33-deficient mice to pneumococcal lung infection.

**Figure 3.**
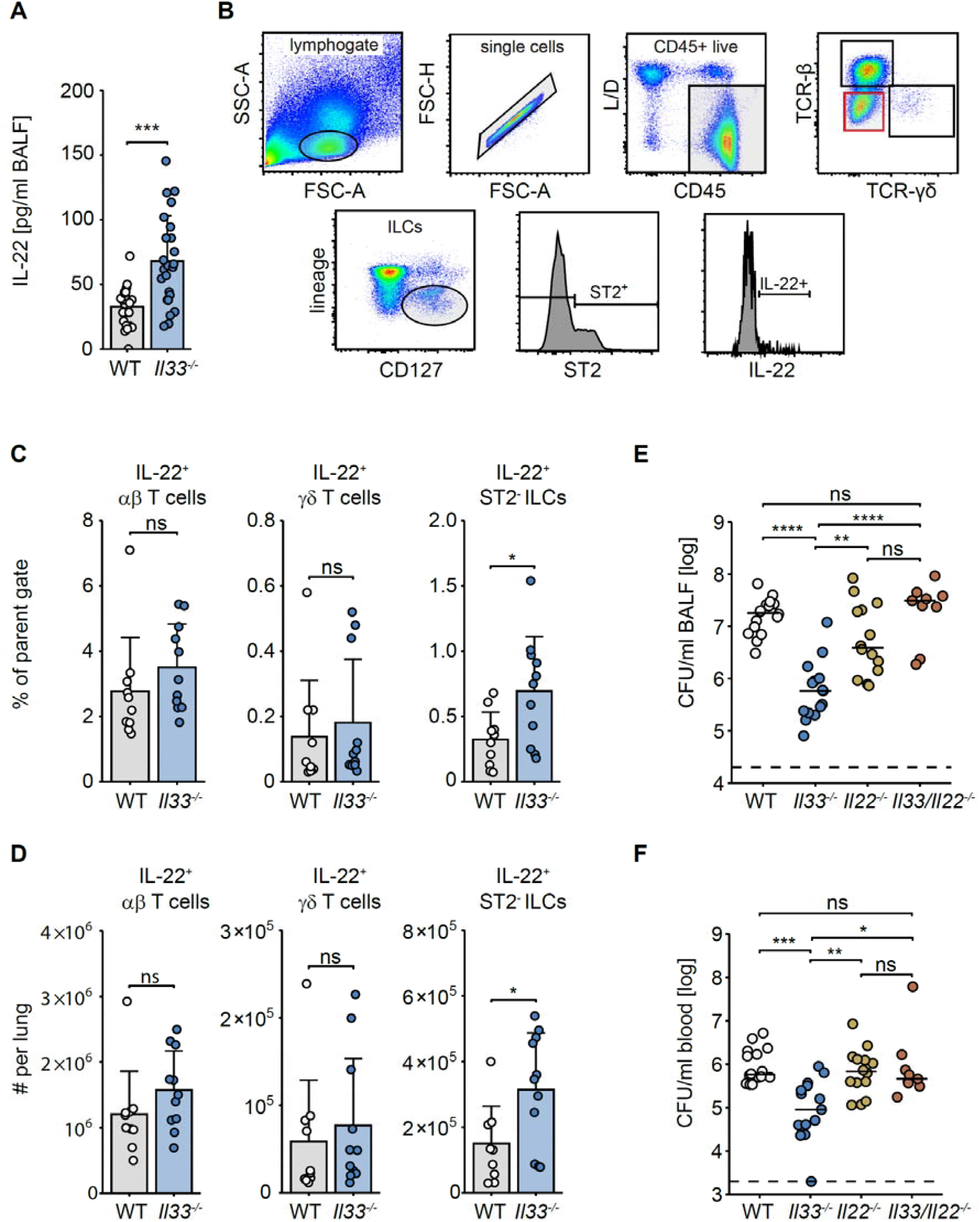
Protective effect of IL-33 deficiency in pneumococcal infection is mediated by enhanced production of IL-22. (**A**) Mice were infected with *S. pneumoniae* and sacrificed after 18 h, and levels of IL-22 in BALF were measured by ELISA, Wilcoxon rank sum test; *** = p < 0.001. (**B**) Representative gating strategy for analyzing IL-22^+^ lymphoid cells by FACS. (**C, D**) Frequency and absolute numbers of IL-22 producing lymphoid cells in lungs of mice 18 h after infection. (**E, F**) WT, *Il33*^-/-^, *Il22*^-/-^ and *Il33*^-/-^*Il22*^-/-^ mice were infected for 48 h and bacterial loads were determined in BALF (**E**) and blood (**F**). Data are shown as individual data points. Bars represent mean + SD, lines represent median and dashed line the lower detection limit, Kruskal-Wallis test followed by Dunn’s posthoc test, ns = p > 0.05, ** = p < 0.01, *** = p < 0.001, **** = p < 0.0001.

### The effect of IL33 deficiency on pneumococcal infection depends on the vivarium in which the mice were bred

For operational reasons, the breeding of our *Il33*^-/-^ and WT control animals had to move several times from one vivarium to another within different institutional breeding facilities in Berlin. Unexpectedly, we observed that the phenotype of the *Il33*^-/-^ mice in *S. pneumoniae* infection differed between the facilities: *Il33*^-/-^ animals from several different vivaria with lower hygienic conditions (e.g. conventional caging instead of individually ventilated cages and/or free access for scientists instead of access only for the responsible animal caretakers) retained their enhanced resistance to pneumococcal infection as compared to WT animals from the same vivaria. However, *Il33*^-/-^ mice from other vivaria with higher hygienic status (vivaria II, IV, V), were less resistant to the infection and thus behaved similarly to WT animals (Fig. 4A, Fig. S5). In contrast, susceptibility of WT mice to pneumococcal infection seemed not to be noticeably influenced by the different housing environments (Fig. S5). These results led us to hypothesize that inter-facility microbiome variability resulted in alterations in the gut microbiota of primarily the *Il33*^-/-^ mice, leading to enhanced defense against *S. pneumoniae* infection in these animals.

**Figure 4.**
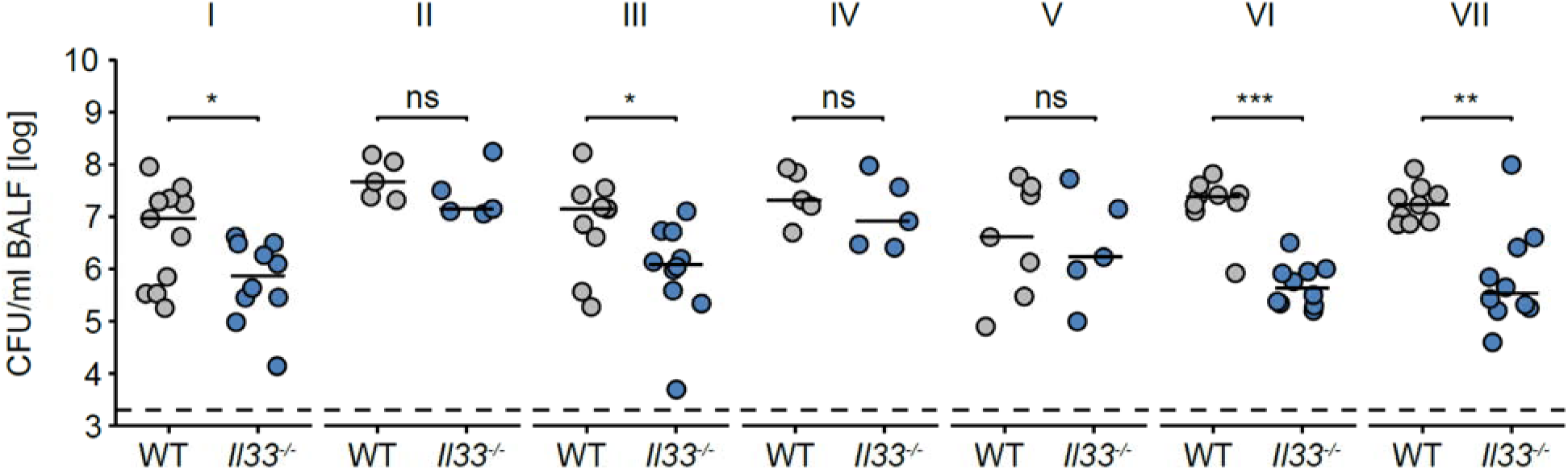
*Il33*^-/-^ from different vivaria differ in their susceptibility to *S. pneumoniae* infection. (**A**) WT and *Il33*^-/-^ mice from various animal vivaria numbered I-VII were infected, sacrificed after 48 h and bacterial loads in BALF were assessed. Data are shown as individual data points. Lines represent median and dashed line the lower detection limit. Wilcoxon rank sum test, * = p < 0.05, ** = p < 0.01, *** = p < 0.001.

### IL-33’s influence on antibacterial defense depends on its modulatory effect on the microbiota

The microbiota has recently emerged as an important factor calibrating immune responses and thereby influencing susceptibility to infection (4, 32-37). To investigate if microbiota differences were indeed responsible for the variable resistance of *Il33*^-/-^ animals to *S. pneumoniae* infection, we first sequenced fecal samples of WT and *Il33*^-/-^ mice from different vivaria to analyze their microbiota. While alpha diversities were comparable between samples of all WT and *Il33*^-/-^ groups (Fig. 5A), taxonomic profiling revealed that the composition of samples from ‘resistant’ *Il33*^-/-^ animals was different compared to samples from ‘susceptible’ *Il33*^-/-^ mice and both WT controls. For example, the microbiota of ‘resistant’ *Il33*^-/-^ mice were characterized by higher relative abundance of various *Lactobacillus* spp. and lower abundance of *Saccharicrinis fermentans* and *Peptococcus niger* as compared to WT animals and to *Il33*^-/-^ animals that did not show the resistant phenotype (Fig. 5B). Moreover, ‘resistant’ *Il33*^-/-^ mice also differed in terms of their gut virome, archean and eukaryotic communities from the other animal groups (Fig. S6A-C). Real-time PCR-based quantification of segmented filamentous bacteria (SFB), which are known to enhance IL-22 production, revealed no difference between WT and *Il33*^-/-^ animals (38, 39), although the relative abundance of SFB in ‘resistant’ *Il33*^-/-^ mice was slightly higher compared to ‘susceptible’ *Il33*^-/-^ animals (Fig. S6D). Importantly, when WT and *Il33*^-/-^ mice from a vivarium normally breeding ‘resistant’ *Il33*^-/-^ animals were treated with antibiotics to deplete their microbiota, the difference in controlling *S. pneumoniae* infection and IL-22 production vanished (Fig. 5C, D). Similarly, cohousing of WT and *Il33*^-/-^ mice normalized their intestinal microbial communities (Fig. 5E, Fig. S6A-C) and equalized their antibacterial defense (Fig. 5F, G). Together, our results demonstrate that an altered microbiota in *Il33*^-/-^ animals is responsible for their enhanced antibacterial resistance during *S. pneumoniae* infection.

**Figure 5.**
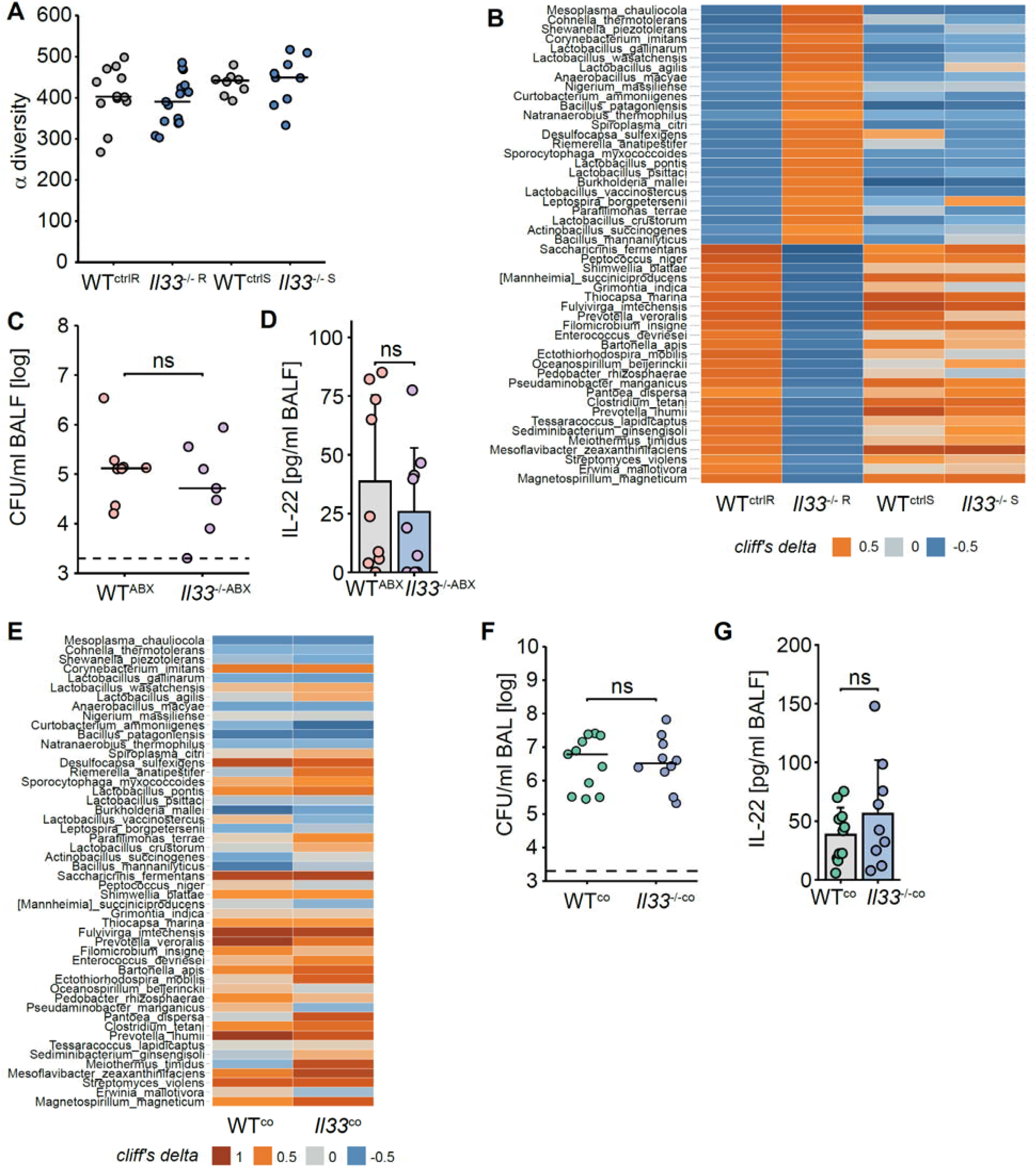
IL-33’s influence on antibacterial defense depends on its modulatory effect on the microbiota. (**A**) Alpha diversity of microbiota samples from *Il33*^-/-^ animals with ‘resistant’ (*Il33*^-/-R^) and ‘susceptible’ phenotype (*Il33*^-/-R^) and corresponding controls (WT^ctrlR^ and WT^ctrlS^) is shown. (**B**) Heatmap of shotgun sequenced microbiota derived from resistant and susceptible WT and *Il33*^-/-^ mice. Cliff’s delta was applied to visualize differences between *Il33*^-/-R^ and WT^ctrlR^. (**C, D**) WT and *Il33*^-/-^ mice were microbiota-depleted by oral treatment with antibiotics,subsequently infected with 1×10^6^ CFU/mouse and sacrificed after 48 h. Bacterial loads (**C)** and IL-22 levels (**D**) were assessed in BALF. Data are shown as individual data points, lines represent median, dashed line the lower detection limit and bars represent mean + SD. Wilcoxon rank sum test, ns = p > 0.05. (**E**) Heatmap of differential microbiota derived from “resistant” and co-housed WT and *Il33*^-/-^ mice. Cliff’s delta was applied to quantify differences (same species as in (**B**)). (**F, G**) WT and *Il33*^-/-^ mice were co-housed for 4 weeks and infected for 48 h. Bacterial loads (**F)** and IL-22 levels (**G**) were assessed in BALF. Data are shown as individulal points, lines represent median, dashed line the lower detection limit and bars represent mean + SD. Wilcoxon rank sum test, ns = p > 0.05.

## DISCUSSION

In our study, we describe a mechanism of bidirectional interaction between the immune system and the microbiota. The mechanism depends on both genetic and environmental factors and influences the efficacy of antibacterial immune defense and thus susceptibility to pneumonia. Specifically, we demonstrate that IL-33 regulates IL-22-dependent antibacterial defense in the lungs. This regulation of antibacterial defense is dependent on IL-33’s modulatory influence on the microbiota and is only apparent in mice that are kept in vivaria with lower hygienic conditions. Mice lacking ILC2s or IL-4Rα did not differ from their controls in response to *S. pneumoniae* infection, although all of them were housed in the same vivarium as the ‘resistant’ *Il33*^-/-^ animals. We speculate that one or several as yet unidentified microorganism(s), which we believe is present only in the microbiotas of mice from low hygienic vivaria, is normally restricted by IL-33 but blooms in *Il33*^-/-^ animals to increase IL-22 production. SFB would have been a candidate for such a microorganism, as it is well known to enhance IL-22 production (38, 39). However, we did not detect increased abundance of SFB in our *Il33*^-/-^ animals as compared to WT controls, thus making it unlikely that these bacteria are causally linked to the enhanced IL-22 production in our *Il33*^-/-^ animals. To narrow down the identity of such a microbe, we compared the microbiota of ‘resistant’ and ‘susceptible’ IL-33-deficient mice and WT controls. Unfortunately, however, the number of taxa that differed in abundance was too large to test them individually in, for example, transplantation experiments to evaluate their ability to induce IL-22-dependent defense against *S. pneumoniae*.

The ‘resistant’ phenotype of *Il33*^-/-^ mice related to the altered microbiota was only observed when the animals were bred in vivaria with lower hygienic standards. Importantly, microbiota alterations in *Il33*^-/-^ mice have also been described by others (40), suggesting that this is not a phenomenon observed specifically in our facility. The previous study, however, found increased abundance of *Akkermansia muciniphila* and SFB in *Il33*^-/-^ mice, whereas our *Il33*^-/-^ mice differ in other taxa from their controls. Overall, we consider our results relevant since animals with lower hygienic status are likely to better reflect the physiological reality. This assumption is consistent with recent studies demonstrating that "wildlings", as mice whose hygiene status could be considered particularly low, mimic human immune responses better than SPF animals kept under high hygienic conditions (41, 42). Moreover, our analyses of a large cohort of patients with community-acquired pneumococcal pneumonia and age- and sex-matched controls revealed an association between pneumococcal pneumonia and carriage of SNP alleles previously linked to altered IL-33 signaling (22-24). These results provide preliminary evidence that IL-33 signaling may also influence susceptibility to bacterial lung infections in humans.

Two elegant studies have recently demonstrated that lung epithelial IL-33 imprints an anti-inflammatory “M2-like” phenotype in alveolar macrophages by acting through ILC2s, basophils and IL-13 (43, 44). In addition, IL-13-deficient mice were shown to exhibit enhanced resistance to bacterial infection (43). In contrast to these studies, we did not observe decreased susceptibility of mice lacking the alpha chain of the IL-4 receptor, which is required for IL-4 and IL-13 signaling. We speculate that differences in the infection models and/or the microbiome of the animals used could explain this discrepancy.

Overall, our study provides novel insights into the interaction between the microbiota and the immune system and how this affects susceptibility to pulmonary infection. Future work is needed to identify the specific microbe(s) which is normally controlled by IL-33 and regulates IL-22 production. Moreover, the mechanism of how IL-33 influences the intestinal microbiota is still unknown and warrants further investigations.

## MATERIALS AND METHODS

### Study approval

All animal experiments were carried out in adherence to the German Animal Welfare Act (Tierschutzgesetz, TierSchG) and to the Federation of European Laboratory Animal Science Association (FELASA) guidelines, following approval by the responsible institutional (Charité – Universitätsmedizin Berlin) and governmental animal welfare authorities (LAGeSo Berlin, approval IDs G0266/11, G0365/12, G0227/16, G0080/18 and T0014/12). Samples from the pneumococcal pneumonia patients were provided by the CAPNETZ foundation (45). This prospective multicenter study (German Clinical Trials Register: DRKS00005274) was approved by the ethical review board of each participating clinical center (Reference number of leading Ethics Committee “Medical Faculty of Otto-von-Guericke-University in Magdeburg”: 104/01 and “Medical School Hannover”: 301/2008) and was performed in accordance with the Declaration of Helsinki. All patients provided written informed consent prior to enrolment in the study.

### Mice

*Il33^-/-^* mice (46) (www.cdb.riken.jp/arg/mutant%20mice%20list.htm; Acc. No. CDB0631K), kindly provided by Dr. Hirohisa Saito (National Research Institute for Child Health and Development, Tokyo, Japan) were crossed with *Il22^-/-^*^(47)^ mice to generate a double knockout line *Il33^-/-^Il22^-/-^*. *Nmur1* ^iCre-eGFP^*Id2* ^fl/fl^ mice and *Id2* ^fl/fl^ littermate controls were recently described ^(28, 48)^. *Il4ra^-/-^* mice were described in Barner et al. (49). *Il1rl1*^-/-^ mice ^(50)^ were kindly provided by Dr. Andrew McKenzie, MRC Laboratory of Molecular Biology, Cambridge, UK. *P2rx7*^-/-^ (51) and *P2ry6*^-/-^ (52) mice were described before and were kindly provided by Dr. Marco Idzko (University Hospital Vienna AKH, Medical University of Vienna, Vienna, Austria). All mice were on a C57BL/6 background and were bred in the animal facility of the Forschungseinrichtungen für experimentelle Medizin (FEM), Charité – Universitätsmedizin Berlin. *Il22^-/-^*, *Il33^-/-^Il22^-/-^*, *Nmur1* ^iCre-eGFP^*Id2* ^fl/fl^, *Id2* ^fl/fl^, *Il1rl1*^-/-^ and *Il4ra^-/-^* mice were bred in the same vivaria that generated ‘resistant’ *Il33*^-/-^ animals. For all experiments female age-matched mice, usually aged 8-12 weeks were used.

### S. pneumoniae *infection*

The murine in vivo model of pneumococcal pneumonia has previously described (53, 54). Briefly, we used the serotype 3 (ST3) strain of *Streptococcus pneumoniae* PN36 (NCTC7978), kindly provided by Prof. Sven Hammerschidt (Ernst-Moritz-Arndt-Universität Greifswald, Germany). Bacteria were plated on columbia agar plates (5 % sheep blood), harvested after 9 h of incubation at 37 °C + 5 % CO_2_, single colonies were picked, optical density was measured at OD_600nm_ and adjusted for the infection dose. Mice were anesthetized with an intraperitoneal (i.p.) injection of 80 mg/kg ketamine and 25 mg/kg xylazine and transnasally inoculated with 5×10^6^ colony forming units (CFU) *S. pneumoniae* in 20 µl PBS per mouse. Sham-infected mice received 20 µl PBS. Following infection, temperature and weight were measured every 12 h to monitor the clinical course of disease. At the indicated time points, mice received an i.p. injection of 160 mg/kg ketamine and 75 mg/kg xylazine and were sacrificed 12, 18, 36 or 48 h post infection by final blood withdrawal. Blood was centrifuged, serum was collected, and lungs were perfused with 5 ml of PBS. All organs of interest were either snap frozen in liquid nitrogen or processed immediately. Bronchoalveolar lavage (BAL) was performed by lavaging the lungs twice with 800 µl PBS containing protease inhibitors. Both fractions were centrifuged and BALF was collected. The pelleted cells were combined and used for subsequent flow cytometry analyses. The bacterial loads were determined in BALF before centrifugation blood and spleen by plating them on blood agar plates and overnight incubation.

### Generation of chimera mice

Mice (10-12 weeks old) were radiated with a dose of 10 Gy and 24 h later transplanted with 5×10^6^ freshly isolated bone marrow cells of either WT or *Il33*^-/-^ congenic (CD45.1^+^) mice. Donor cells were first reconstituted in 200 µl PBS and then intravenously injected into recipient mice. To prevent infections, recipient mice received 0.01 % enrofloxacin *ad libidum* for four weeks starting one day before radiation. After 10 weeks of reconstitution, mice were infected with *S. pneumoniae* and bacterial defense was investigated.

### In vitro infection

Bone marrow-derived macrophages (BMM), alveolar epithelial cells (AEC) or whole lung suspensions were isolated from WT mice, infected in vitro with 1×10^6^ CFU *S. pneumoniae* (D39) and supernatant was harvested 16 h (BMM and AEC) and 24 h (murine lung homogenate) post infection. Protein levels of uric acid, ATP and IL-33 were measured via ELISA (eBioscience, R&D Systems) according to the manufacturer’s instructions.

### Intratracheal treatment of mice

WT mice were intratracheally treated with PBS (control), uricase (0.073 U/per mouse), apyrase (0.32 U/per mouse), suramin (32 μM/per mouse) or pyridoxal-phosphate-6-azophenyl-2‘,4‘-disulfonic acid (PPADS) (32 μM/per mouse) dissolved in 25 μl PBS immediately before *S. pneumoniae* infection and 24 h after. For the treatment 24 h post infection, mice were orotracheally intubated with a laryngoscope and the solution was then applied into the lungs with a microsprayer. Mice were sacrificed after 48 h and bacterial burden was quantified.

### Real-time qPCR

DNA extraction was carried out utilizing the zymoBIOMICS DNA extraction kit (Zymo), following the instructions provided by the manufacturer. Gene expression analysis was conducted using 1x SYBR Green Master Mix, following the manufacturer’s guidelines from Applied Biosystems. The primer sequences for qRT-PCR were the following: SFB_forward: 5’-GACGCTGAGGCATGAGAGCAT-3’, SFB_reverse: 5’-GACGGCACGGATTGTTATTCA-3’.

### Flow cytometry

The cell pellets obtained from both lavages were pooled in 1 ml PBS and analyzed by flow cytometry. The relative and absolute number of alveolar macrophages (CD45^+^ CD11b^-^ Siglec-F^+^), recruited polymorphonuclear neutrophils (CD45^+^ CD11b^+^ Ly6G^+^) and inflammatory monocytes (CD45^+^ Ly6G^-^ CD11b^+^ Ly6C^++^) were determined. Moreover, lungs were perfused, isolated, cut into pieces and incubated in 5 ml RPMI 1640 (+ 5 % FCS) containing collagenase (150 U/ml) and DNase (1500 U /ml) for 40 min at 37 °C while shaking. Digested lung cells were passed through a 70 µm cell strainer and red blood cells were lysed. Cells were solved in FACS-buffer (PBS + 2 % FCS) and subsequently stained for flow cytometry. Following populations were measured in lungs: Innate lymphoid cells (ILC) (CD45^+^ CD127^+^ TCRβ^-^ TCRγδ^-^ CD3^-^ CD19^-^ FcεRIα^-^ Ly6G^-^ CD5^-^ NK1.1^-^ Siglec-F^-^), innate lymphoid cells type II (ILC2) (CD45^+^ CD127^+^ CD90^+^ CD3^-^ CD19^-^ FcεRIα ^-^ Ly6G^-^ CD5^-^ NK1.1^-^ GATA3^+^), αβ T cells (CD45^+^ CD3^+^ TCRβ^+^ TCRγδ^-^) and γδ T cells (CD45^+^ CD3^+^ TCRβ^-^ TCRγδ^+^). For intracellular staining, lungs were prepared as described, left unstimulated or stimulated with 50 ng/ml Phorbol-12-myristat-13-acetat (PMA), 1 µg/ml Ionomycin and 1 µg/ml Brefeldin A (all Sigma-Aldrich) for 5 h at 37 °C + 5 % CO_2_. For the process of permeabilization either FoxP3 Fix/Perm kit (Biolegend) or 2% paraformaldehyde (PFA) was used. ILC2 were sorted for CD45^+^ CD3^-^ CD19^-^ Ly6G^-^ FceRIa^-^ CD5^-^ NK1.1^-^ CD127^+^ CD90^+^ ST2^+^ and further processed for bulk RNA sequencing.

### ELISA

Quantification of cytokine and chemokine levels in BALF was conducted by using commercial a multiplex assay (ProcartaPlex, Thermo Fisher) and commercially available single cytokine ELISA kits (eBioscience, R&D Systems).

### Single cell RNA sequencing

After infection and preparation, lungs were perfused with 5 ml PBS followed by 2 ml dispase (5000U/ml, Corning). 700 µl dispase was applied intratracheally, followed by 500 µl lukewarm 1 % low melt agarose. Lungs were then removed and transferred into digestion medium (PBS + 5 % FCS + 2 µg/ml Actinomycin D + 2 mg/ml collagenase + 0.5 mg/ml DNase), cut into pieces and incubated for 30 min at 37 °C at 125 rpm in a shaker. Lung homogenates of each group were combined, filtered through a 70 µm cell strainer, red blood cells were lysed, and cells were counted using a hemocytometer and dead cells were removed with a magnetic bead based dead cell removal kit according to the manufacturer’s instructions (EasySep STEMCELL technologies). Following lung digestion 1/4 of the cell suspension was sorted for CD45^+^ CD3^-^ CD19^-^ Ly6G^-^ FceRIa^-^ CD5^-^ NK1.1^-^ CD127^+^ CD90^+^ ILCs and spiked back into the whole lung single cell solution. Then cells were loaded into the Chromium Controller (10x). Single Cell 5’ reagent kit v2 was used for reverse transcription, cDNA amplification and library construction, followed by the detailed protocol provided by 10x Genomics. The generated libraries were sequence on a NovaSeq 6000 S4 flowcell type with 200 cycles with 30,000 cells per lane and 4 lanes in total, targeting 2,500 lanes/lane. The reads were aligned to the reference genome provided by 10x Genomics (Mouse reference mm10), and a digital gene expression matrix was generated to record the number of unique molecular identifiers (UMIs) for each gene in every cell. Quality metrics were applied to the count matrices, and thresholds were set for the number of genes (> 150 and < 4000) and the percentage of mitochondrial reads (less than 10%). For data integration, variable feature finding, data scaling, and PCA calculation based on highly variable genes, the standard pipeline of Seurat package (version 4.2.0) was used. In case of batch effects, the Harmony package (55) or SoupX (56) was utilized for correction. The functions RunUMAP(), FindNeighbours() and FindClusters() were employed to reduce dimensions and find Shared Nearest Neighbours (SNN) in the dataset. To calculate differential gene expression, the Seurat function FindMarkers() was used.

### Fecal DNA extraction and shotgun sequencing

Mouse fecal samples derived from mice bred in vivaria II, III, IV, V, VI and VII were collected before infection and stored at -80 °C until further use. For the DNA extraction a DNA Miniprep Kit from Zymo Research (#D4300) was used, purity and concentration of the DNA was measured using a NanoDrop^TM^ 2000 (Thermo-Fisher). Library prep and sequencing of the bacterial DNA was performed by Eurofins Genomics (Ebersberg, Germany) using an Illumina NovaSeq 6000, with 10 million reads and paired end sequencing (2 x 150bp). After sequencing, preprocessing was conducted by Eurofins genomics. Poor quality bases, adapters and primers were removed before proceeding to the removal of host sequences. Taxonomic profiling was conducted by using the NCBI database of bacterial, archaeal, fungal, protozoan and viral genomes. FASTQ files were generated and a table containing normalized reads for each domain, phylum, class, order, family, genus and species was provided. This table was then loaded into R and used for subsequent analyses.

### Microbiota depletion

To study the effects of antibiotic treatment on the gut microbiota of mice, 8-9-week-old mice were housed in sterile cages and treated orally with a combination of imipenem (250 mg/L; Fresenius Kabi), metronidazole (1 g/L; Braun), vancomycin (500 mg/L; HIKMA Pharma), ciprofloxacin (200 mg/L; Fresenius Kabi) and ampicillin (1 g/L; Ratiopharm) in the drinking water *ad libitum* for a period of 6 weeks (ABX mice) as described earlier (57). Fecal samples were collected weekly for analysis of fungal outgrowth and depletion of the gut microbiota, using bacterial and fungal cultivation on agar plates. Any mice that showed fungal outgrowth were excluded from the study.

### Bulk RNA sequencing

To prepare mouse lungs for RNA bulk sequencing, infected lungs were digested and FACS-sorted for ILC2 as described. RNA was isolated using Direct-Zol Microprep Kit (Zymo Research, Cat#R2061). Library preparation was done with Takara SMARTer stranded Total RNA-Seq Kit v3 - Pico Input and library was sequenced on a NovaSeq 6000 SP targeting 400 million reads. Adapter sequences were removed from sequencing reads with Cutadapt (v3.7) and their quality assessed with FASTQC (v0.11.9). Subsequently, reads were input into SeA-SnaP: (Se)q (A)nalysis (Sna)kemake (P)ipeline (https://github.com/bihealth/seasnap-pipeline). Briefly, reads were mapped with STAR aligner (v2.7.3a) to the mm10/GRCm38 mouse genome using GENCODE annotation version M12. Next, featureCounts (v2.0.0) was used to count reads in exons and generate a read count table. Differential expression analysis was carried out with DeSeq2 (v1.34.0) ^56^ Differentially expressed genes (adjusted P value <0.1) were subjected to gene set enrichment analysis (GSEA) using the fgsea package (version version 1.16.0) ^56^.

### SNP analysis

Samples were provided by the CAPNETZ competence network, a German multi-center prospective cohort study for CAP (45). The control groups consisted of a subgroup of healthy adults of similar age and sex distribution from the PolSenior program, an interdisciplinary project, designed to evaluate health and socio-economic status of the Polish Caucasians aged ≥65 y (58). DNA Genotyping of rs1420101, rs7044343, rs9500880 and rs1921622 was conducted by PCR utilizing fluorescent-labeled hybridization FRET probes followed by melting curve analysis in a LightCycler 480 (Roche Diagnostics). The following probes and primers were used: rs1420101: f-primer: TAgTTTggTgTCAgAgTTTCTgCAA, r-primer: TgAAgTgACTTACTCAAggCCA, anchor probe: LC640 CCAATgAgTATTACTAAAgATTAAgCTCTT-PH, sensor probe: AAAgCCTCTCATTAAACTTTgAA-FL; rs7044343: f-primer: AggAATgAATATTgggTgACACTATg, r-primer: TACCCAAgTTCAAgAggCACTg, anchor probe: LC640 TCCTgTCTgCATgTAAAgCCACTC-PH, sensor probe: ggTTACTTCTCAgggCATCA-FL; rs9500880: f-primer: TTCAggAAATTAgAggTCTAATgTAA, r-primer: CTgCAgAgACATgCCAAAgACA, anchor probe: LC640 gACgAAAgCATTCTTAAATCTgATATTC-PH, sensor probe: TgATTTCTAgTTCCACACTTATgA-FL; rs1921622: f-primer: CACCAggATAACTCTgCCAC, r-primer: TAAATTTgCAAATgTTTCACCAAC, anchor probe: LC640 gCCATAggCACTAgCTgAAATAC-PH, sensor probe: TAAAAATTgATgAATTTTgTTCTgg-FL.

### Data analysis and statistics

Data analysis was performed using R, version 4.2.1. Wilcoxon rank-sum test was used for the comparison of two groups. If more than two groups were compared, Kruskal-Wallis Test, followed by a Dunn’s post-hoc test was utilized. SNP allele frequencies were analyzed using Fisher’s exact test.

## ACKNOWLEDGEMENTS

The authors are grateful to all patients for consenting to biosampling and data collection. We would like to thank Ulrike Fiebiger, Soraya Mousavi, Katrin Heiden, and the staff of the animal facility of the Forschungseinrichtungen für experimentelle Medizin (FEM) of the Charité – Universitätsmedizin Berlin for excellent technical assistance, animal breeding and generation of secondary abiotic mice. We also would like to thank Dr. Katrin Reppe for her help with our animal experiments, and Dr. Lutz Hamann for his advise regarding SNP analysis.

## SUPPLEMENTARY FIGURES

**Figure S1.**
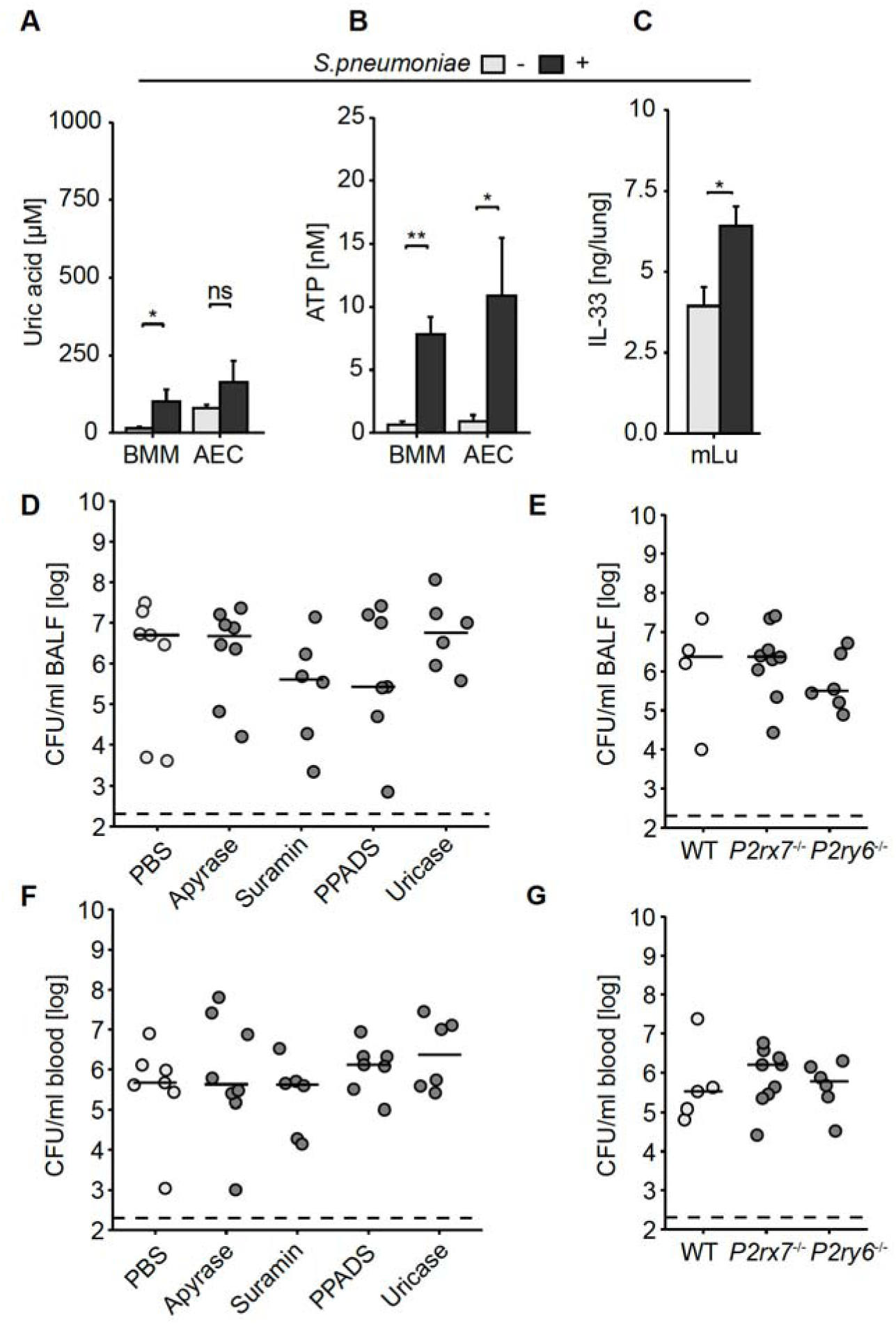
Role of alarmins in *S. pneumoniae* infection. Murine bone marrow-derived macrophages (BMM) and alveolar epithelial cells (AEC) and murine lung tissue (mLu) were left untreated or infected with 10^6^ CFU of *S. pneumoniae* for 16-24 h. (**A**) Uric acid, (**B**) ATP and (**C**) IL-33 were measured. Data are shown as mean + SEM; n = 8-14 per group; Wilcoxon rank sum test; ns = p > 0.05, * = p < 0.05, ** = p < 0.01. (**D-G)** C57BL/6J WT mice were left untreated or treated with different alarmin inhibitors and WT, *P2rx7*^-/-^ and *P2rx6*^-/-^ mice were infected with *S. pneumoniae* and bacterial loads in BALF (**D, E**) and blood (**F, G**) were measured. Data are shown as individulal points, lines represent median and dashed lines the lower detection limit.

**Figure S2.**
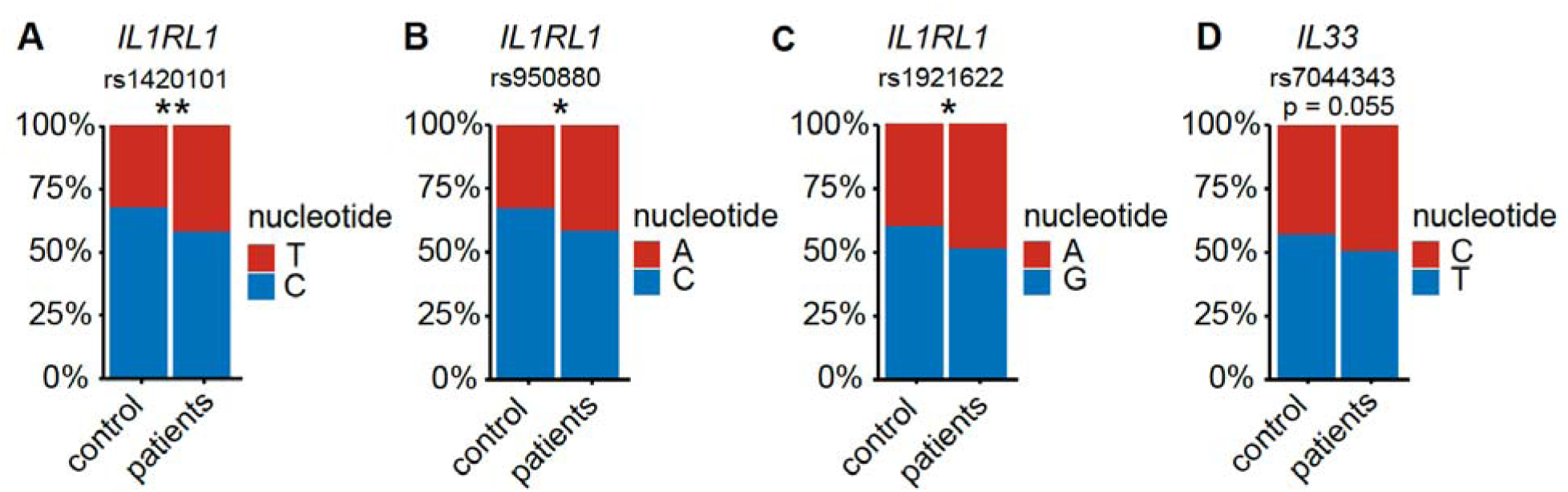
SNP in *IL33* and *IL1RL1* are associated with pneumococcal pneumonia. (**A-D**) Frequencies of SNP alleles of the *IL33* and *IL1RL1* genes were assessed in 238 patients with community-acquired pneumococcal pneumonia patients and 238 age- and sex-matched controls. Allele frequencies are visualized; Fisher’s exact test. .** = p < 0.01, * = p < 0.05, ns = p > 0.05.

**Figure S3.**
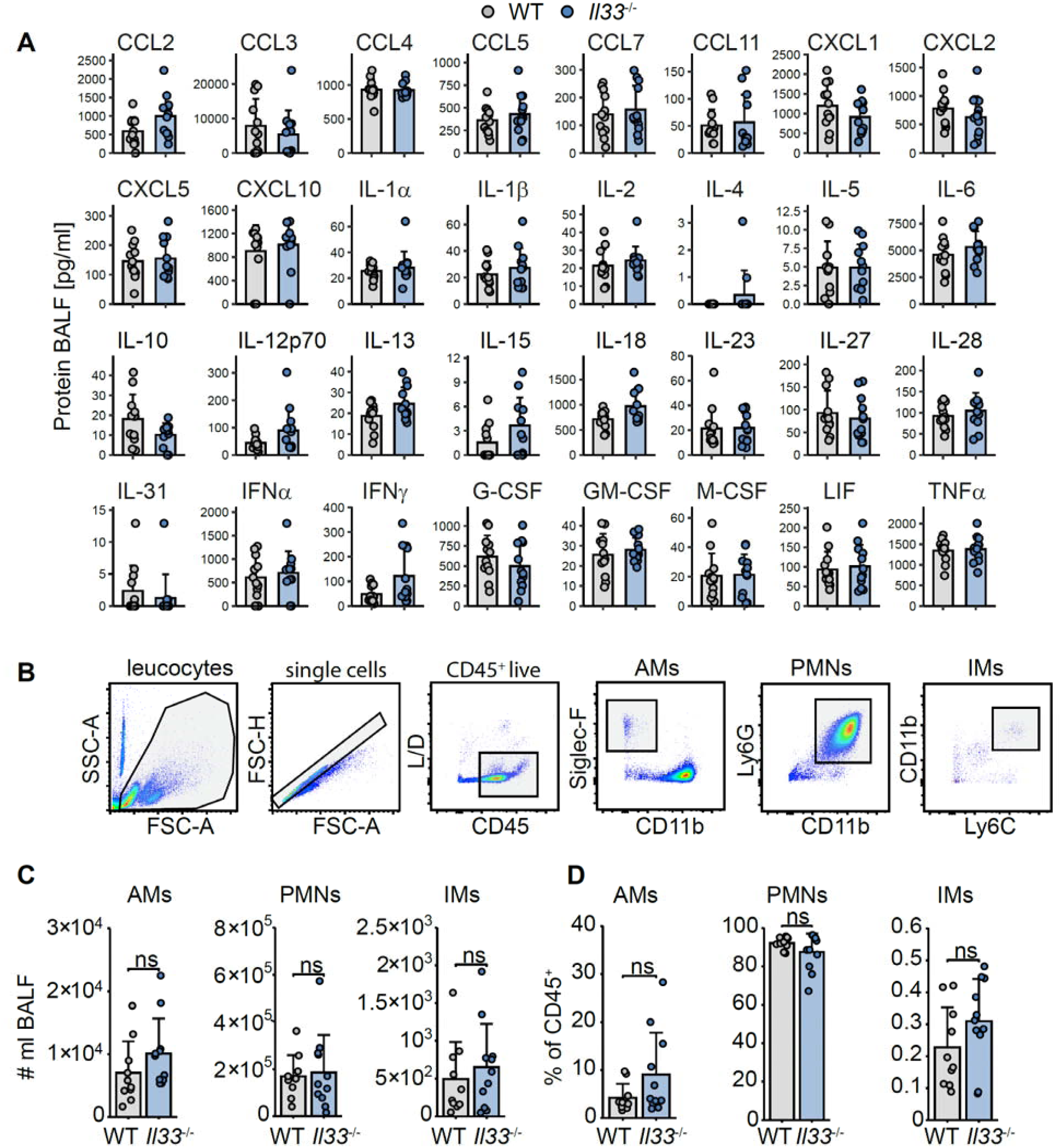
IL-33 deficiency does not influence production of several inflammatory cytokines or number or proportion of major innate leukocytes. (**A**) WT and Il33^-/-^ mice were intranasally infected with *S. pneumoniae*. After 18 h, cytokine and chemokine levels in BALF were quantified by multiplex ELISA. (**B**) Representative gating strategy to analyze macrophages, PMNs and inflammatory monocytes (IMs) by flow cytometry. (**C, D**) WT and *Il33^-/-^* mice were infected, sacrificed after 18 h, and absolute numbers and frequencies of leucocytes were measured in BALF by flow cytometry. Bars represent mean + s.d., Wilcoxon rank sum test; ns = p > 0.05.

**Figure S4.**
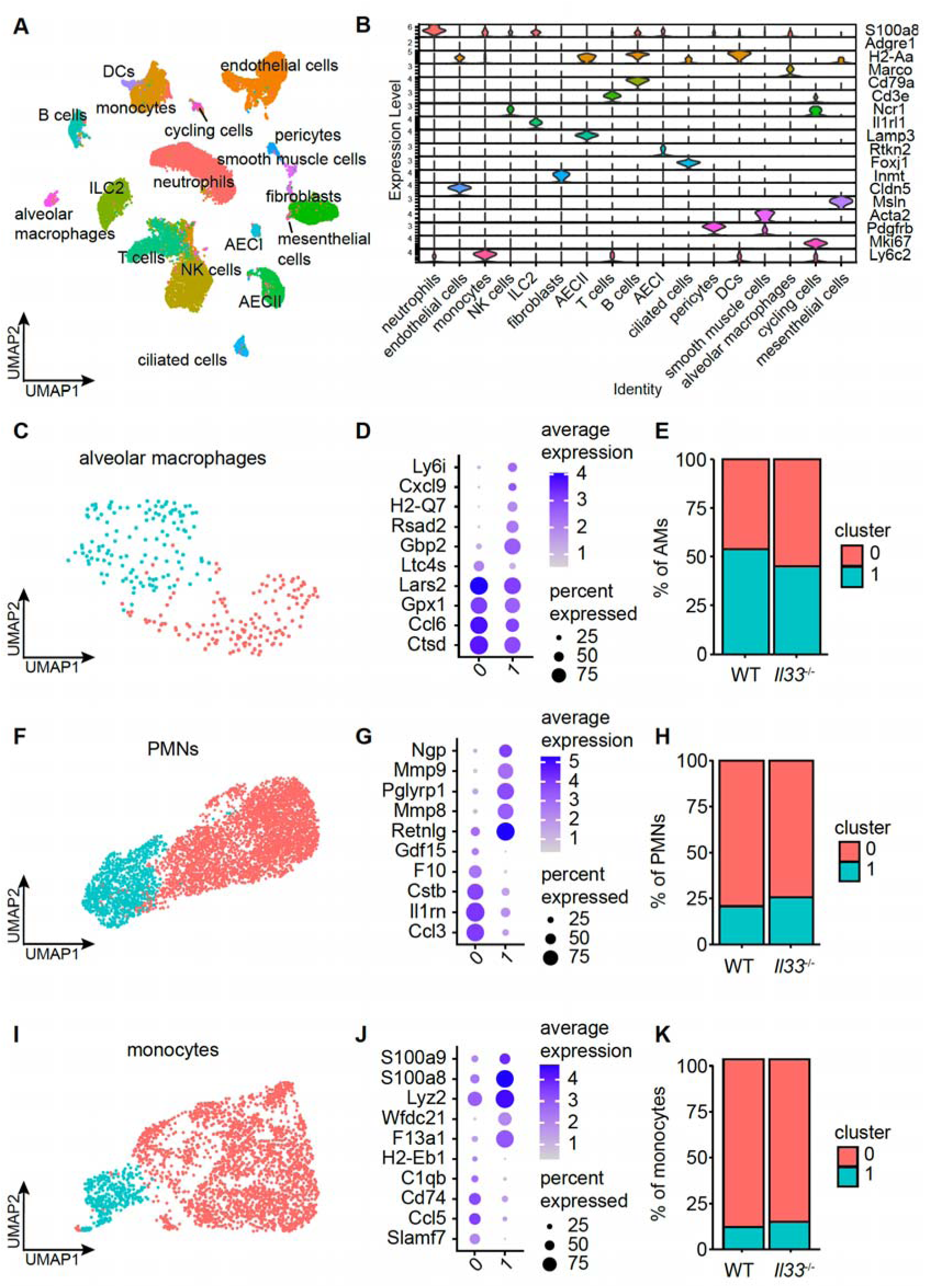
scRNAseq analysis of pulmonary cells from WT and *Il33*^-/-^ mice. Mice were infected, sacrificed after 36 h (n = 3-4 per group) and lungs were subjected to scRNAseq. (**A**) Two-dimensional embedding computed by UMAP on 24612 computationally identified cells. (**B**) Stacked violin plot depicting representative marker genes for each cell type. (**C – K**) Dataset was first subsetted on alveolar macrophages, PMNs and monocytes and separated in two clusters by unbiased clustering (**C, F, I**). Dotplots of cluster specific marker genes (**D, G, J**) and frequencies in WT and *Il33*^-/-^ are represented in a barplot (**E, H, K**).

**Figure S5.**
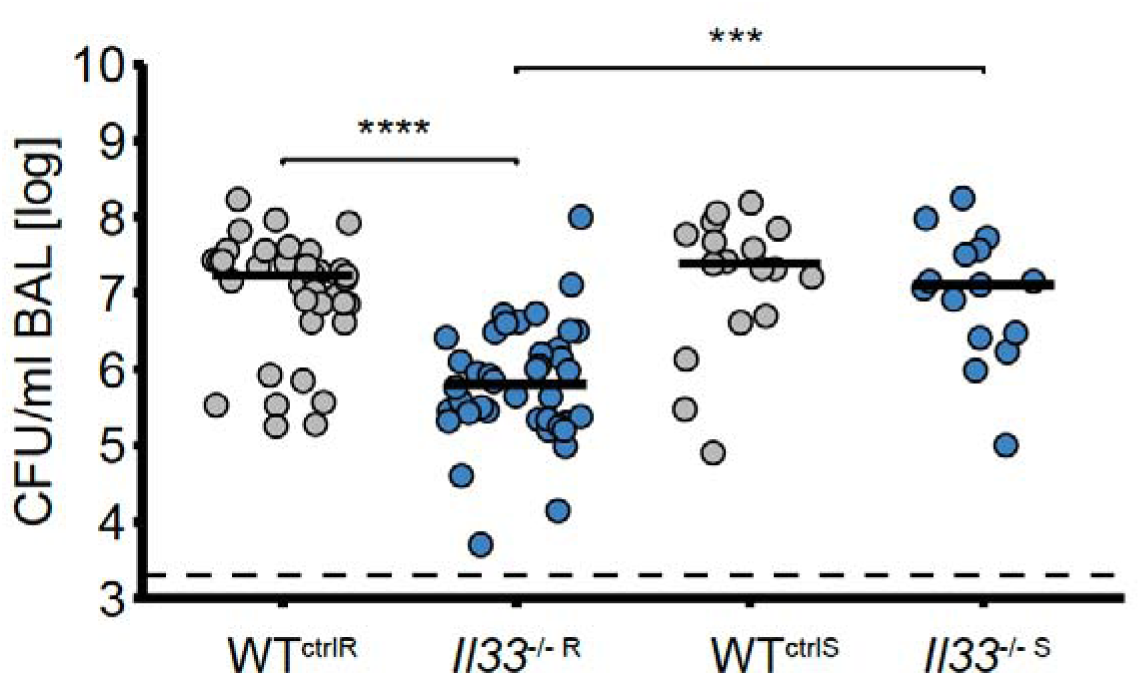
Susceptibility to *S. pneumoniae* infection differs in *Il33*^-/-^ but not WT mice from different vivaria. Accumulated CFU data from Fig. 4 classified by genotype and phenotype. (‘resistant’ *Il33*^-/-^ = *Il33*^-/-R^, corresponding WT = WT^ctrlR^, ‘susceptible’ *Il33*^-/-^ = *Il33*^-/-S^, corresponding WT = WT^ctrlS^). Data are shown as individual data points, lines represent median and dashed line lower detection limit, Kruskal-Wallis followed by Dunn’s posthoc test; *** = p < 0.001, **** = p < 0.0001.

**Figure S6.**
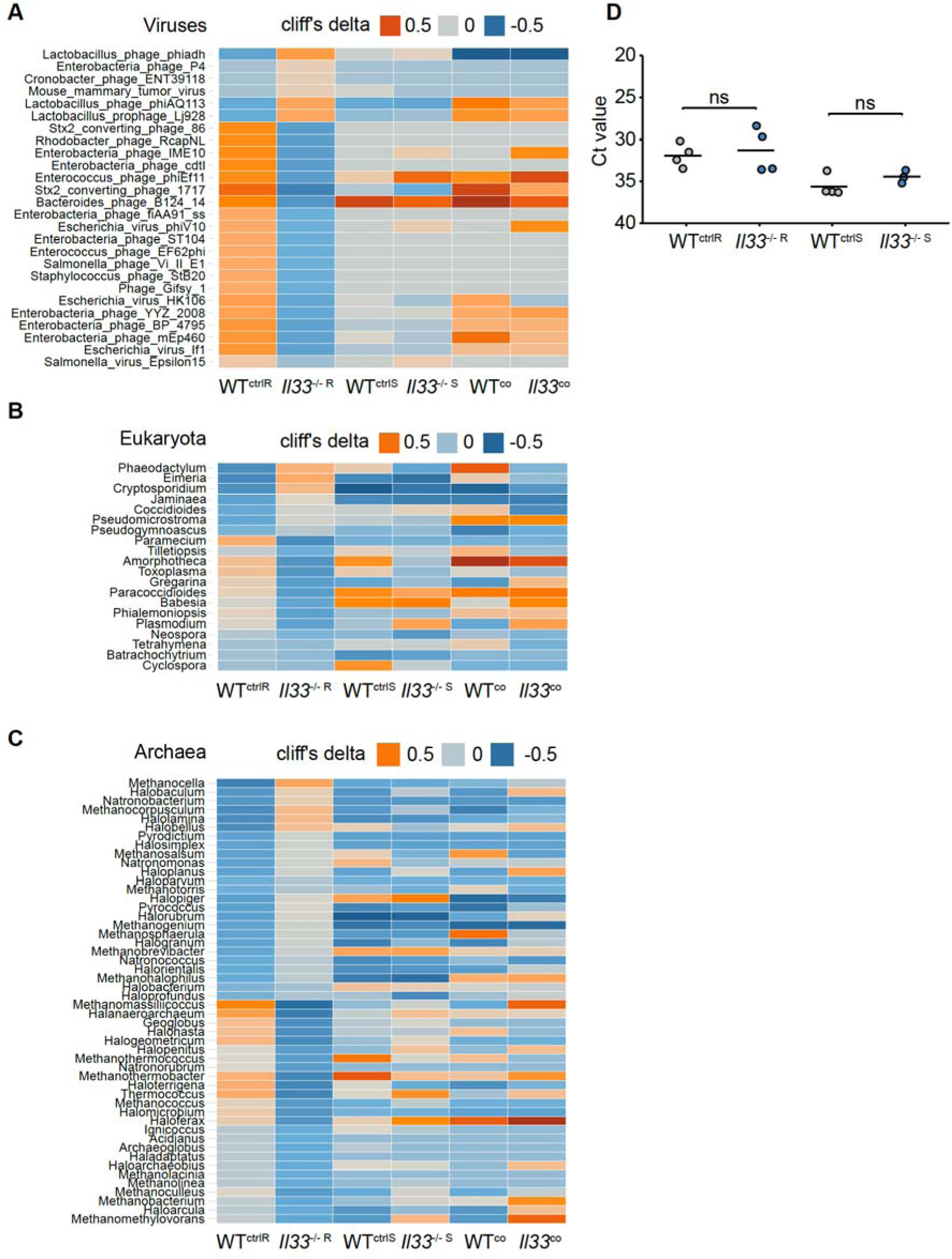
IL-33’s influence on the gut virome and eukaryotic microbial communities. Heatmap of shotgun sequenced microbiota derived from resistant, susceptible and co-housed mice (‘resistant’ *Il33*^-/-^ = *Il33*^-/-R^, corresponding WT = WT^ctrlR^, ‘susceptible’ *Il33*^-/-^ = *Il33*^-/-S^, corresponding WT = WT^ctrlS^). Cliff’s delta was applied to quantify differences in viruses (**A**), eukaryota (genus level) (**B**) and archaea (genus level) (**C**) between *Il33*^-/-R^ and WT^ctrlR^ mice. (**D**) SFB was quantified in fecal samples from *Il33*^-/-R^, *Il33*^-/-S^, WT^ctrlR^ and WT^ctrlS^ mice by PCR. Lines represent median. Wilcoxon rank sum test, ns = p > 0.05.

